# Close-Up of vesicular ER Exit Sites by Volume Electron Imaging using FIB-SEM

**DOI:** 10.1101/2025.04.23.650353

**Authors:** Athul Nair, Akhil Nair, Patrick Stock, Arkash Jain, Elliott Somerville, Anwesha Sanyal, Jose Inacio Costa-Filho, Tom Kirchhausen

**Affiliations:** Department of Cell Biology, Harvard Medical School, 200 Longwood Ave, Boston, MA 02115, USA; Program in Cellular and Molecular Medicine, Boston Children’s Hospital, 200 Longwood Ave, Boston, MA 02115, USA; Department of Pediatrics, Harvard Medical School, 200 Longwood Ave, Boston, MA 02115, USA

**Keywords:** endoplasmic reticulum, secretory pathway, COPI, COPII, Deep learning, Automated Segmentation in Electron Microscopy (ASEM)

## Abstract

Volume electron microscopy by high-pressure freeze substitution combined with block-face focused ion beam scanning electron microscopy (FIB-SEM) provides comprehensive 3D views of subcellular architecture, essential for understanding cellular activity in context. Aided by Automated Segmentation in Electron Microscopy (ASEM)—a 3D-Unet convolutional neural network trained with sparse annotations—we characterized the spatial organization of endoplasmic reticulum exit sites (ERES), the initial locations for membrane remodeling in protein secretion, in cells not overexpressing secretory cargo.

The ASEM model, trained on 50–70 nm in diameter COPI vesicles, successfully identified vesicles adjacent to the Golgi apparatus of HeLa, SVG-A, and iPSC-derived neurons. It also revealed abundant vesicles of similar appearance, often within a group of ∼5–40 vesicles clustered within a 250 nm^3^ region adjacent to flattened endoplasmic reticulum (ER) domains, forming what we propose are COPII-mediated vesicular ER-exit sites. Elsewhere, similar assemblies appeared alongside tubular ER networks emerging from similarly enlarged ER domains previously reported as the only ERES in HeLa cells.

These findings underscore the power of large-scale volume electron microscopy to resolve contradictions regarding membrane organization in vesicular trafficking. We encourage the scientific community to use our publicly accessible repository, containing open-source code, trained models, annotations, predictions, and FIB- SEM datasets, to facilitate continued advances in automated segmentation methods.

**Summary:** Nair et. al. show abundance of vesicular ERES in mammalian cells imaged by volume focused ion beam electron microscopy.

## INTRODUCTION

Volume electron microscopy provides unparalleled views into the 3D subcellular organization, offering a wealth of contextual information critical for understanding membrane dynamics. Among these, focused ion beam scanning electron microscopy (FIB-SEM) enables large-scale, three-dimensional imaging of intracellular structures with near-isotropic high resolution. Most FIB-SEM datasets achieve 8-nm resolution (Narayan et al., 2014; Knott et al., 2008; Xu et al., 2017; Wu et al., 2017; Hoffman et al., 2020)), with recent developments reaching 4-5 nm (e.g. (Xu et al., 2021; Müller et al., 2020; Sanyal et al., 2024; Gallusser et al., 2022; Weigel et al., 2021; Heinrich et al., 2021)). When applied to cells preserved by high-pressure freezing and freeze substitution (HPFS), FIB-SEM maintains native ultrastructure while providing a comprehensive view of macromolecular assemblies, membranes, and organelles—within individual cells or even across clusters of cells in intact tissues. This approach has detected spatial relationships that were previously obscured by the limited depth of field in conventional thin-section transmission electron microscopy (TEM), which images 30- to 300-nm- thick sections, representing only a small fraction of the whole cell (e.g., (Ladinsky et al., 1994; Zeuschner et al., 2006; Mironov et al., 2003)).

Despite its potential, the complexity of such FIB-SEM datasets makes direct visual inspection impractical. To address this limitation, AI-based methods powered by sparsely labeled ground truths and a 3D-Unet convolutional neural networks have been developed to automate segmentation, greatly improving the identification and characterization of subcellular structures (Heinrich et al., 2021; Gallusser et al., 2022; Mekuč et al., 2022). The Cell Organelle Segmentation in Electron Microscopy (COSEM) project at the Janelia Research Campus uses deep learning to train models for detecting and quantifying intracellular structures within large-scale EM datasets (Xu et al., 2021). The Automated Segmentation of Intracellular Structures in Electron Microscopy (ASEM) pipeline, independently developed by our group, also uses deep neural networks to identify organelles ranging from clathrin-coated vesicles and nuclear pores to mitochondria, the Golgi apparatus, and the endoplasmic reticulum (Gallusser et al., 2022; Galbraith, 2023). These AI-driven approaches now greatly accelerate the detailed characterization of organelles in both normal (Gallusser et al., 2022; Heinrich et al., 2021) and metabolically altered states (Parlakgül et al., 2022).

Taking advantage of these advances, we investigated a fundamental membrane remodeling event concerning the anterograde trafficking of cargo from the endoplasmic reticulum (ER) to the Golgi apparatus. Trafficking that originates at ER exit sites (ERES) was initially proposed to involve COPII coat assembly at the ER surface with the resulting ∼50–70 nm COPII-coated vesicles then functioning as transport carriers (Balch et al., 1994) [reviewed in (Béthune and Wieland, 2018)]. Support for this model originated from *in vitro* reconstitution assays showing formation of ∼50 nm COPII vesicles from ER membranes of yeast and mammalian cells [reviewed in in (Béthune and Wieland, 2018)]. High-resolution 3D immuno-gold labelling transmission electron microscopy of ∼400 nm sections of frozen human liver derived HepG2 cells provided further *in vivo* morphological evidence, revealing COPII and cargo-containing 50-60 nm vesicles alongside ∼200 nm tubular structures near expanded ER domains (Zeuschner et al., 2006). Cryo-tomography of cryo-FIB milled INS-1 rat insulinoma cells active in secretion showing omega-shaped buds emerging from flattened ER cisternae, closely associated with clusters of ∼52–64 nm coated and uncoated vesicles, as well as small ∼100 nm pearled tubules, but the resolution was insufficient to definitively identify the vesicle coat type (Carter et al., 2020). The tubules might correspond to the previously proposed “mega-carriers” or extended tubular carriers implicated in the ER-to-Golgi transport of large cargoes, such as lipoproteins and ∼300-nm procollagen assemblies (Gorur et al., 2017; Raote and Malhotra, 2019). Observations showing vesicles and pearled-like tubules at the region between the ER and Golgi in fat bodies and imaginal discs of flies were later reported using volume focused ion beam electron microscopy (FIB-SEM) (Yang et al., 2021). A cryo-tomography study of *C. reinhardtii* focused on the Golgi apparatus revealed multiple COPI-coated buds with dense, uniform coats at cisternal rims and clathrin coated vesicles near the trans-Golgi network regions (Bykov et al., 2017). Although not the primary focus, ERES adjacent to the Golgi contained slightly larger COPII-coated vesicles with a characteristic two-layered coat; however, no data were presented from other cellular regions. Taken together, these morphological observations support conclusions suggesting the existence of COPII-coated vesicular carriers, independently reached through numerous biochemical experiments conducted under diverse conditions, including analyses of partially and fully purified membranes and COPII components carriers. These findings contrast with earlier work (Mironov et al., 2003), which used chemically fixed samples under conditions of synchronized ER exit of a large cargo or membrane protein and reported an absence of vesicular intermediates—proposing instead that non-coated tubular carriers emerge directly from the ER.

Recently, concerns were raised about the interpretation of these structural studies supporting vesicle-mediated transport from ERES, citing potential limitations related to section thickness, limited sampling volume, and, in some cases, artifacts introduced by chemical fixation (Weigel et al., 2021). To address these issues, Lippincott-Schwartz and colleagues applied high-resolution volumetric FIB-SEM imaging at isotropic 4–8 nm resolution, using cells preserved by HPFS. They complemented this approach with correlative light-electron microscopy using cryo-structured illumination (Hoffman et al., 2020). Observing cells undergoing synchronized cargo release in cells ectopically expressing cargo or COPII subunits, they concluded that only tubular and not vesicular carriers emerged from enlarged ER domains. They also reported a similar presence of tubular ERES and absence of vesicular ERES in non-transfected Hela cells, challenging the classical model of COPII vesicles as the key transport intermediates for conventional cargo in mammalian cells.

We report here our effort to resolve this apparent contradiction by imaging cells devoid of ectopic expression and preserved by HPFS, using high-resolution volumetric FIB-SEM. We analyzed the 3D images aided by the ASEM pipeline, trained to independently recognize COPI vesicles and endoplasmic reticulum (ER). This approach enabled reliable identification of 50–70 nm COPI-coated vesicles adjacent to the Golgi apparatus, as well as COPI-like vesicles located near the ER and spatially separated from the Golgi. These vesicles typically formed clusters of 3 to 40, adjacent to flattened ER cisternae, were slightly larger, had a different appearance from COPI vesicles, and appeared throughout the cell volume in all mammalian cell types examined. We refer to these assemblies as vesicular ERES.

Considering these observations, we re-examined the same FIB-SEM dataset of a HeLa cell lacking ectopic expression of COPII subunits or secretory cargo (Weigel et al., 2021). Analysis of the automated COSEM segmentation—released to the COSEM repository post-publication—revealed that approximately two-thirds of the structures annotated as tubular ERES contain vesicle-rich regions. Together, these findings support the coexistence of both vesicular and tubular ERES architectures in mammalian cells.

## RESULTS

### FIB-SEM Imaging

We used volume FIB-SEM to image cells preserved by HPFS (Hoffman et al., 2020; Studer et al., 2008; Xu et al., 2021). Our dataset included two human astrocyte-derived SVG-A cells (SVG-A1 and SVG-A2), one immortalized mouse dendritic cell (MutuDC), and a human induced pluripotent stem cell (iPSC)-derived neuron (iN) cultured *in vitro* (Table S1). Volume images were acquired using crossbeam FIB-SEM at an isotropic resolution of 5 × 5 × 5 nm per voxel or anisotropic resolution of 2 x 2 x 1 nm. We also analyzed a volumetric dataset of the HeLa-2 cell from the COSEM project acquired with a custom-modified FIB-SEM at 4 × 4 × 5.24 nm (Weigel et al., 2021) (Table S1). Unless otherwise noted, we trained and applied our ASEM model (Gallusser et al., 2022) directly to raw FIB-SEM volumes and conducted all subsequent visual inspections and analyses on these datasets. In some cases, we examined FIB-SEM datasets acquired at 2 × 2 × 1 nm or 4 × 4 × 5.24 nm resolution denoised with a novel algorithm we developed to be published elsewhere.

### Recognition of COPI Vesicles

COPI vesicles, which measure 50–70 nm in diameter, localize to the periphery of the Golgi apparatus. They mediate selective retrograde transport from the Golgi to the ER and support bidirectional trafficking between Golgi cisternae (Béthune and Wieland, 2018). These vesicles are coated by COPI coatomers but lack a strongly electron-dense coat in FIB-SEM images of HPFS-preserved samples stained with OsO₄ and uranyl acetate. They are nonetheless readily distinguishable from nearby clathrin-coated vesicles, which are slightly larger (70–90 nm) and have a prominent electron-dense coat.

Ground truth segmentations of COPI vesicles were generated by annotating COPI vesicles at the Golgi periphery in FIB-SEM images from SVG-A1 and SVG-A2 cells. We trained the ASEM model (Gallusser et al., 2022) using annotations from 42, 47, and 51 manually segmented COPI vesicles of 50-70 nm in diameter across three distinct regions of interest in SVG-A1, and 32 vesicles from a fourth region in SVG-A2. Annotated volumes ranged from 204 to 250 voxels in x and y axes and 110 to 250 voxels along the z-axis, providing sufficient context for accurate neural network segmentation. An independent set of 35 COPI vesicles within a 180 × 180 × 210 voxel region of SVG-A1 excluded from training was reserved for validation, as detailed in the Methods section. Annotations were performed on datasets acquired at 5 nm resolution using Neuroglancer, a WebGL-based volumetric data viewer (https://github.com/google/neuroglancer) and WebKnossos (Boergens et al., 2017).

The ASEM COPI model achieved a Dice coefficient of 0.8482 during training and 0.4176 for validation, with a final loss (binary cross entropy) of 0.0095 after 167,000 epochs (∼21 hours of training). Visual inspection confirmed accurate identification of COPI vesicles adjacent to the Golgi apparatus in the training and validation datasets, as well as other vesicles in additional regions not included in training of the SVG-A, MutuDC, iN and HeLa-2 cells (Figs. 1-5).

**Figure 1.**
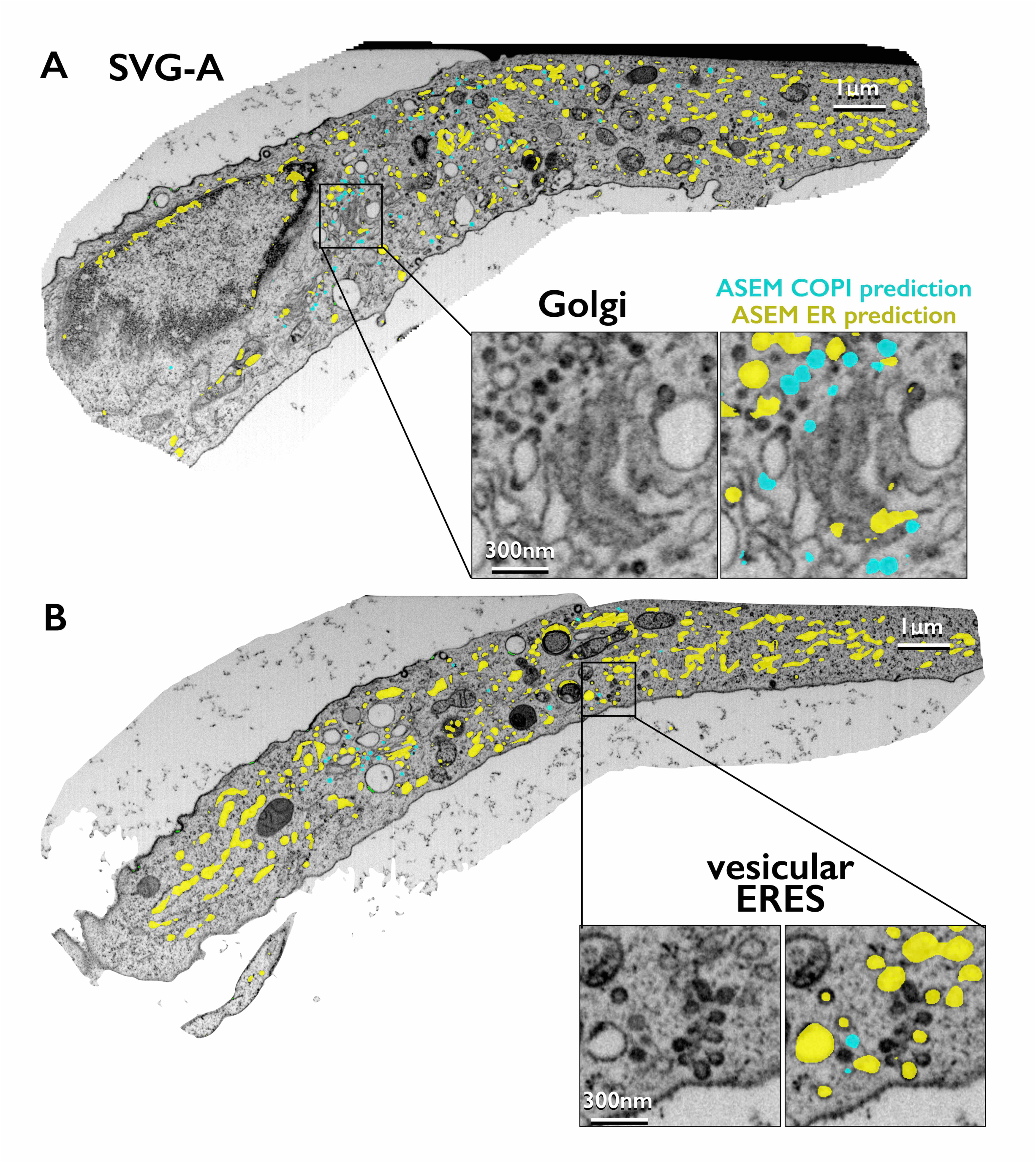
FIB-SEM images and deep-learning neural network ASEM Predictions of ER and COPI vesicles. **(A,B)** Single-plane views from a FIB-SEM volume image of a portion from an interphase SVG-A cell prepared by HPFS and visualized at 5-nm isotropic resolution. Binary masks generated by the ASEM models for ER and COPI highlight predicted structures: yellow for the ER, light blue for COPI vesicles highlighting examples at a location near the Golgi apparatus corresponding to bona fide COP-I vesicles **(A)** and at location adjacent to the ER for COPI-like vesicles assigned as part of the vesicular ERES **(B)**. Scale bars: 1 µm and 300 nm in insets.

A second class of slightly larger vesicles, approximately 90-110 nm in diameter, characterized by a distinct electron-dense coat typical of clathrin-coated vesicles and localized to the distal side of the trans-Golgi network (TGN), was used as an internal control to test the specificity of the COPI model. While these vesicles were not recognized by our COPI ASEM model (Fig. 2), they were accurately identified by our clathrin-coat ASEM model (Fig. 2), previously trained using ground truth annotations of endocytic clathrin-coated pits in plasma membrane images (Gallusser et al., 2022); likewise, the clathrin-coat ASEM model did not misidentify COPI vesicles (Fig. 2). These results illustrate the specificity and robustness of our COPI and clathrin coat models, underscoring the importance of tailored training for specific vesicle classes.

**Figure 2.**
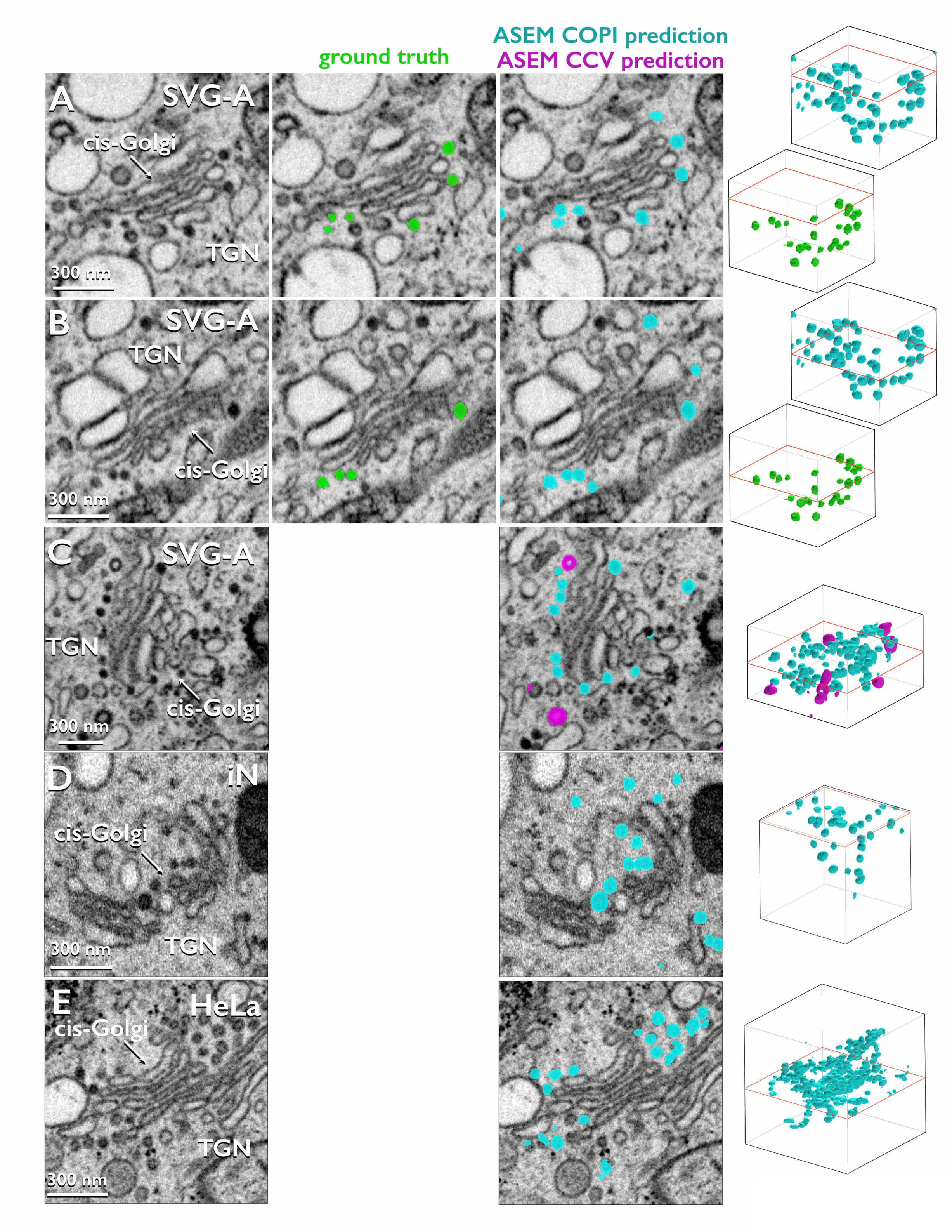
ASEM Predictions of COPI vesicles and Clathrin-Coated Vesicles Near the Golgi Apparatus (Associated to Videos 1-4) **(A-E)** Left panels: Single-plane views from FIB-SEM volume images of the indicated cell types, prepared by HPFS and visualized during interphase at 5-nm isotropic resolution. Central panels: examples of ground truths used for training **(A)** and validation **(B)** for COPI vesicles (green) near the Golgi apparatus imaged at different planes of the same cell. Right panels, predictions near the Golgi apparatus for COPI vesicles (light blue) and clathrin-coated vesicles (CCV, magenta), generated by two different ASEM models trained with COPI and endocytic clathrin coated pits, respectively. Scale bars: 300 nm. Right most panels: Volumetric renditions of the ground truths and ASEM predictions of COPI and clathrin coated vesicles (CCV).

### Identification of Vesicular Endoplasmic Reticulum Exit Sites (ERES)

Additional vesicles, also measuring approximately 50–90 nm in diameter, were identified by our ASEM COPI model across the cell volumes of SVG-A, iN, and HeLa cells. Initial visual inspection using Neuroglancer showed that these vesicles were often located far from the Golgi apparatus (Fig. 1). They were typically members of small clusters of similar vesicles, of which only one or a few had been detected by ASEM. We initially identified approximately 400 such sites in the partial volumes of SVG-A and iN cells and across a fraction of the full volume of the non-transfected HeLa-2 cell.

The high spatial density of organelles and vesicles—ranging from small carriers to larger structures such as endosomes, mitochondria, and ER—complicated the assignment of specific identities to the mapped vesicles. Thus, to aid in the visual interpretation and improve contextual characterization of the vesicles, we took advantage of the extensive distribution of the ER as a three-dimensional spatial reference. Using our previously trained and validated ASEM ER segmentation model (Gallusser et al., 2022), we identified ER within the same cells. These vesicles, recognized by our ASEM COPI model and located away from the Golgi apparatus, were typically within ∼120 nm of flattened, expanded ER cisternae. Detailed visual inspection of a random subset of 30 sites in SVG-A cells and 43 sites in HeLa cells revealed considerable variability in vesicle cluster sizes, ranging from 5 to 40 vesicles per cluster within volumes approximately 400 nm in all directions (Figs. 3, 6). Vesicle diameters measured near the Golgi in SVG-A (59.2 ± 9.1 nm, n = 24) and HeLa (60.1 ± 8.0 nm, n = 24) cells were significantly smaller (p < 0.0001) than those near the ER in SVG-A (75.7 ± 9.1 nm, n = 24) and HeLa (75.9 ± 9.5 nm, n = 24) cells. These dimensions agree with prior measurements of the smaller COPI and larger COPII vesicles.

**Figure 3.**
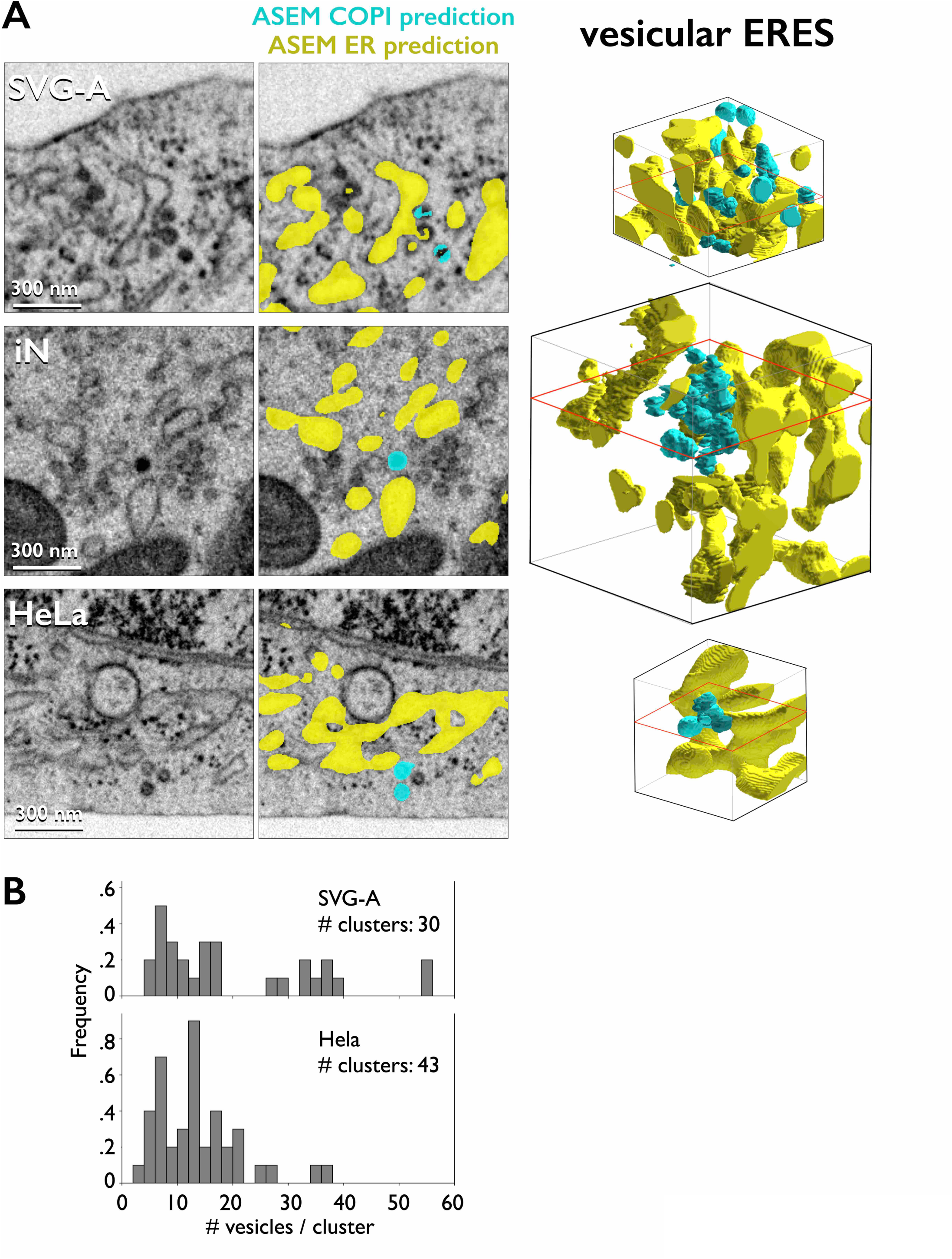
vesicular ERES (Associated to Videos 5-7) Left panels: Single-plane views from FIB-SEM volume images of the indicated cell types, prepared by HPFS and visualized during interphase at 5-nm isotropic resolution. Middle panels: ASEM model predictions of COPI- like vesicles (light blue) clustered near the predicted ER (yellow). Scale bars: 300 nm. Right panels: Manually annotated volumetric renditions of vesicles and ER in vesicular ERES.

Since COPI and COPII vesicles differ in coat composition and architecture, the ability to distinguish them directly in FIB-SEM images would be a significant advantage. However, prior studies using voxel sizes of 4–5 nm have not achieved this. We resolved this limitation by combining FIB-SEM acquisition at 2 × 2 × 1 nm resolution with our novel denoising protocol (to be described elsewhere). Visual inspection of datasets from SVG-A (Fig. 4) and MuTuDC (Fig. 5) cells revealed differences between the smaller, densely stained COPI-coated vesicles near the Golgi and the larger vesicles with a clearer outline of the surrounded membrane associated with ribosome-free ER at ERES (blue arrow heads). The overall staining in the MuTuDC sample was denser and the contrast in appearance between COPI and ER-associated vesicles was less pronounced (Fig. 5). We include an example from the SVG-A cell of vesicles in a closely apposed, concatenated arrangement which, if not well resolved, could be misinterpreted as a continuous tubule (Fig. 4B, C).

**Figure 4.**
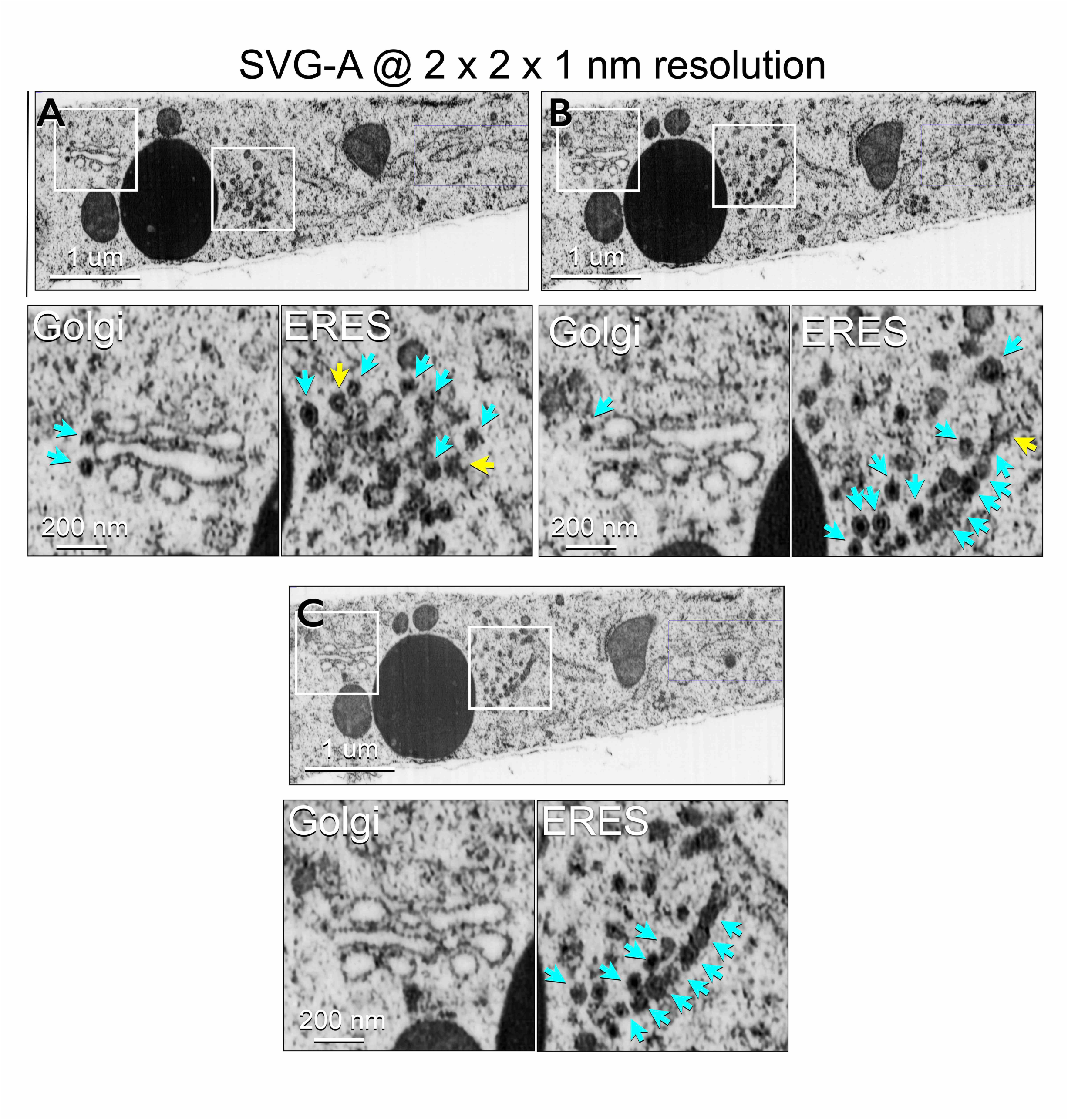
COPI vesicles Near the Golgi Apparatus and vesicular ERES (Associated to Video 8) Single-plane views from a denoised FIB-SEM volume of an interphase SVG-A cell acquired at 2 × 2 × 1 nm resolution. Scale bar: 1 µm. Insets (3× enlargement, scale bar: 200 nm) show light blue arrowheads marking visually confirmed isolated COPI vesicles and associated ERES; yellow arrowheads indicate buds emanating from the ER. **(A-C)** correspond to z planes 384, 524 and 539, respectively.

**Figure 5.**
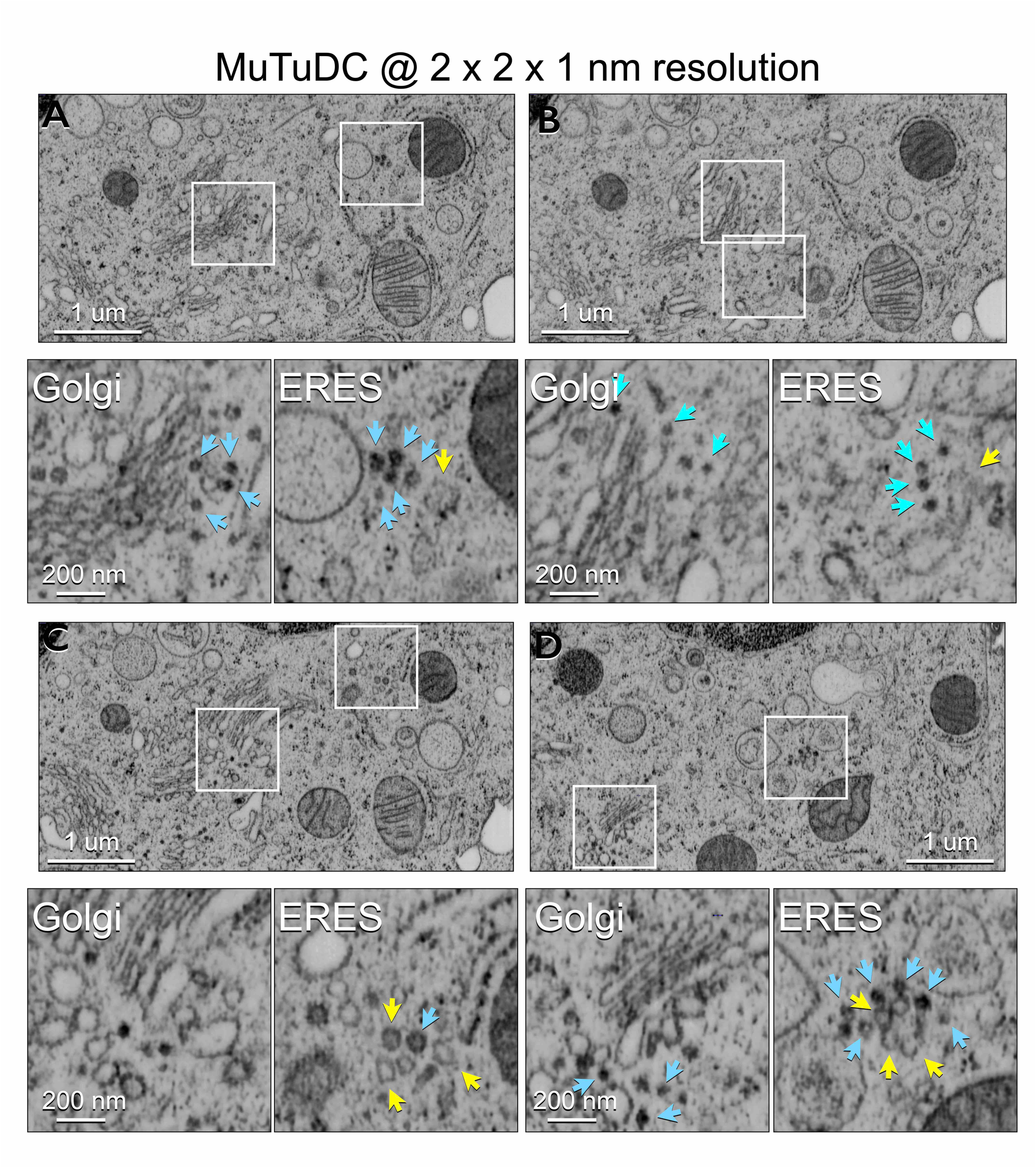
COPI vesicles Near the Golgi Apparatus and vesicular ERES (Associated to Video 9) Single-plane views from a denoised FIB-SEM volume of an interphase MuTuDC cell acquired at 2 × 2 × 1 nm resolution. Scale bar: 1 µm. Insets (3× enlargement, scale bar: 200 nm) show light blue arrowheads marking visually confirmed isolated COPI vesicles and associated ERES; yellow arrowheads indicate buds emanating from the ER. **(A-D)** correspond to z planes 332, 497, 664 and 1552, respectively.

While we acknowledge that the current FIB-SEM data do not provide sufficient structural detail to directly assign molecular identity to the ER-associated vesicles, the available evidence supports a parsimonious interpretation. Given that COPII is the sole known coat for anterograde carriers emerging from the ER, we associate the vesicles observed at ER exit sites with COPII, without inferring a specific state within the coating or uncoating cycle.

We further compared the positions 247 vesicular ERES we identified in the HeLa-2 cell guided by the location of vesicles identified to be in proximity of the ER with 779 tubular ERES annotated in the COSEM project dataset data acquired at 4 x 4 x 5.4 nm resolution (Weigel et al., 2021), which employed an automated tubular ERES segmentation model not used in the original publication (Fig. 6). The vesicular ERES showed no preferential distribution within the cell volume (Fig. 6B). Among these, 179 vesicular ERES, containing vesicle clusters, were spatially distinct from 705 tubular ERES, highlighting heterogeneity in ERES organization. The remaining 68 vesicular ERES, each associated with 1–3 vesicles, localized adjacent to 74 tubular ERES, including instances where two vesicular ERES neighbored a single tubular ERES (Fig. 6A-C).

**Figure 6.**
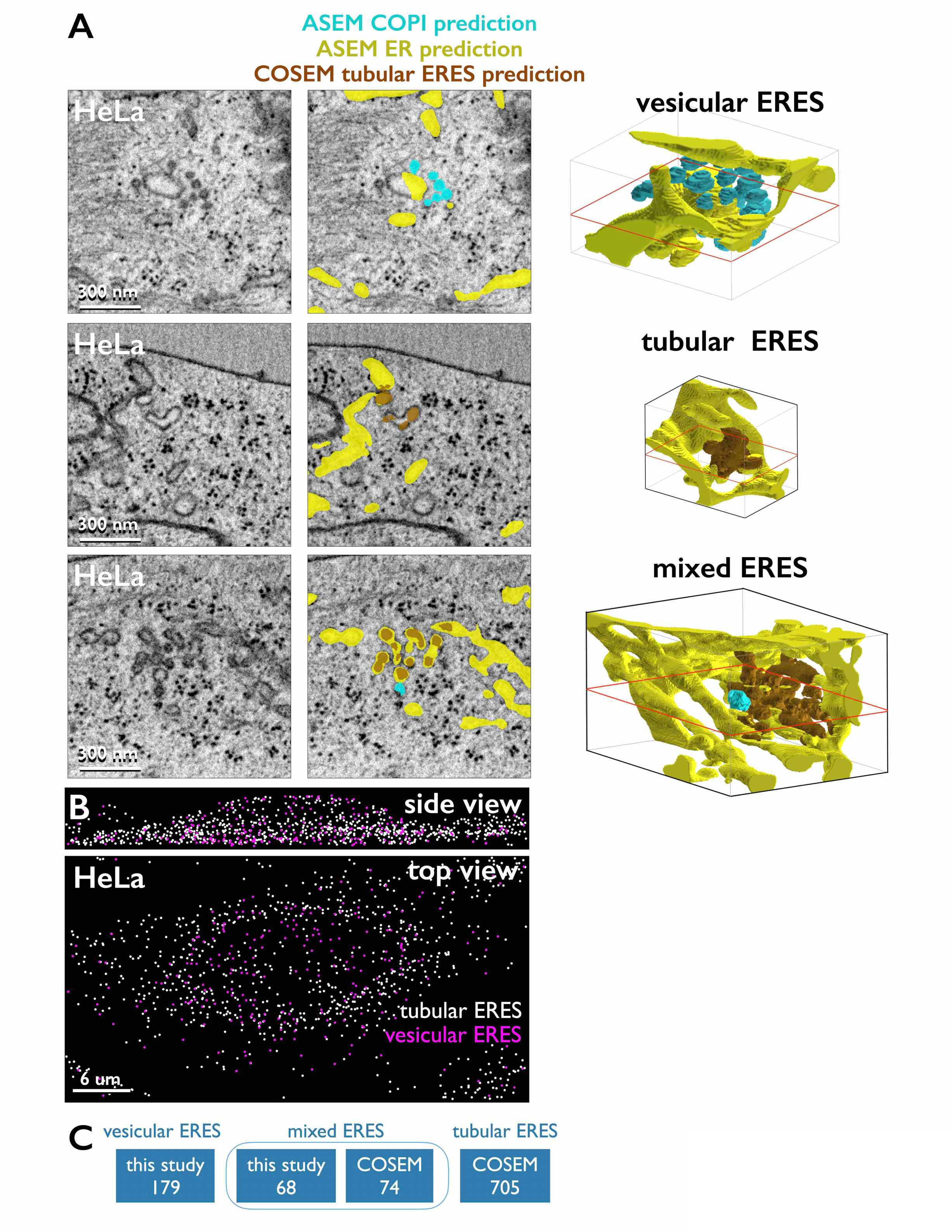
Comparison of Predictions of Vesicular and Tubular ERES (Associated to Videos 10-12) FIB-SEM dataset at 4 × 4 × 5.24 nm resolution generated by the COSEM is from Hela-2 cell during interphase prepared by HPFS. **(A)** Left panels: Single-plane views highlight representative examples of stand-alone vesicular and tubular ERES, and a case where both types are juxtaposed. Middle panels: Predictions of vesicles (light blue) and adjacent ER (yellow) from ASEM COPI and ER models, and tubular membranes (dark orange) from the COSEM model. Scale bars: 300 nm. Right panels: Manually annotated volumetric renditions of vesicles and ER in single vesicular and tubular ERES and in a region containing both types. **(B)** Top and bottom panels: Z-projection views of centroid positions for vesicular (cyan) and tubular (white) ERES. Vesicular ERES (n = 247) were identified using COPI and ER ASEM models; tubular ERES (n = 779) were annotated from the same HeLa-2 cell using the tubular ERES COSEM model. **(C)** Summary of spatial relationships between vesicular and tubular ERES shown in **(B)**. Among the 247 vesicular ERES, 179—each containing clusters of 5–50 vesicles—were spatially distinct from the 779 tubular ERES and are referred to as stand-alone vesicular ERES. The remaining 68 vesicular ERES, each containing 1–3 vesicles, overlapped with 74 tubular ERES and are referred to as mixed ERES. No isolated vesicles were associated with the other 705 tubular ERES.

Finally, we used Neuroglancer to reexamine random regions of the same HeLa-2 cell, denoised from the original 4 × 4 × 5.4 nm dataset, focusing on sites annotated as tubular ERES by the model from (Weigel et al., 2021). Many annotations corresponded to enlarged membrane varicosities, generic ER clusters, or vesicles (Fig. 7), with spatial arrangements resembling the vesicular ERES observed in SVGA, iNs, and MuTuDC cells. Notably, vesicle clusters appeared at approximately two-thirds of the sites annotated as tubular ERES.

**Figure 7.**
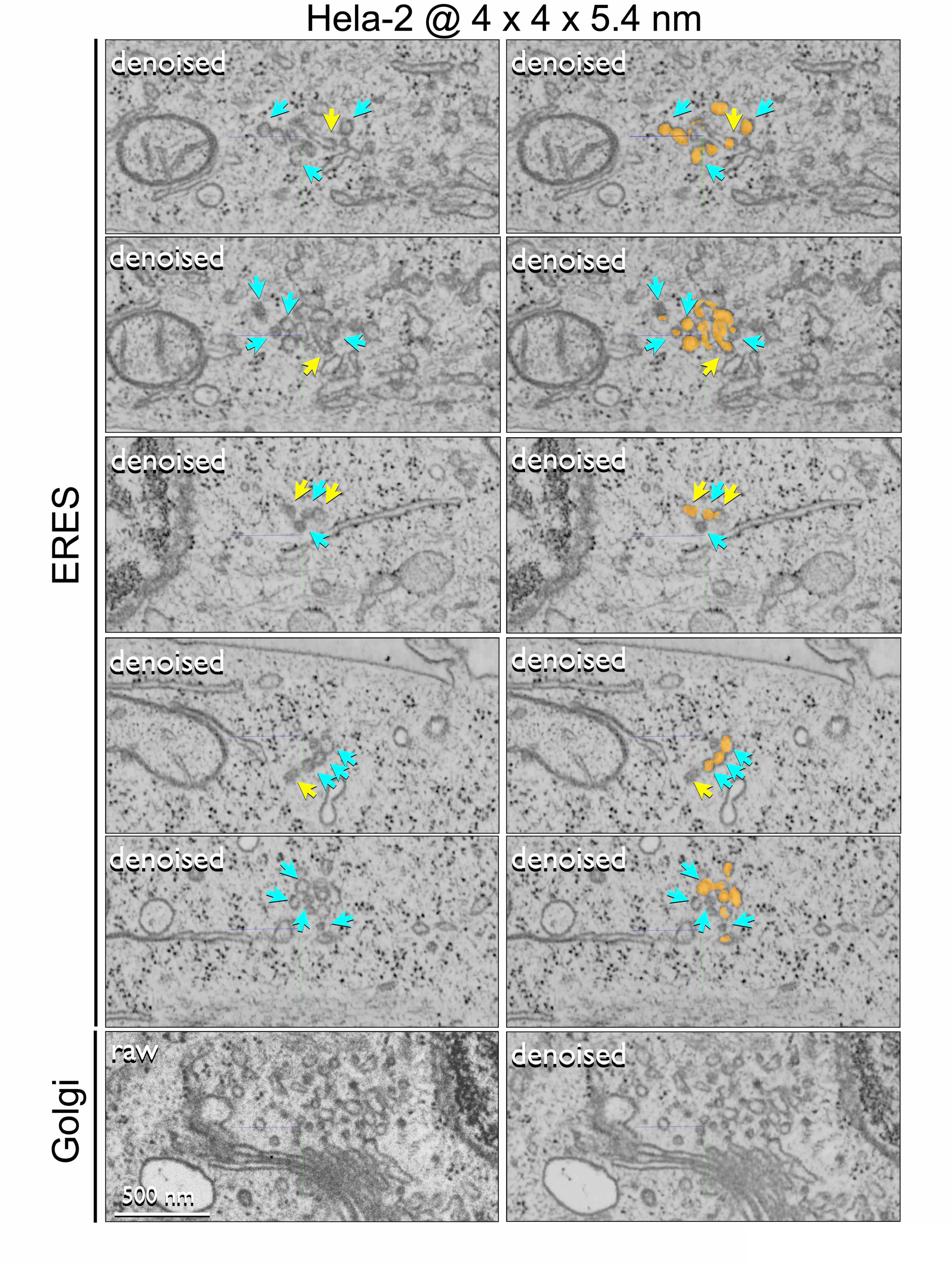
Abundant vesicles at tubular ERES predicted by the COSEM project. Single-plane views from a denoised FIB-SEM dataset of an interphase HeLa-2 cell prepared by HPFS, acquired at 4 × 4 × 5.24 nm resolution. Images show randomly selected sites predicted as tubular ERES by the COSEM model (orange), highlighting abundant isolated vesicles (blue arrowheads) adjacent to tubular ER (yellow arrowheads), verified by visual inspection using Neuroglancer. A representative example near the Golgi apparatus comparing the original and denoised images is also shown. Plane coordinates (z, y, x) in nanometers for the images, from top to bottom, are: (12,506, 5,348, 77,460), (12,452, 5,348, 37,460), (15,090, 4,028, 16,276), (17,093, 4,059, 13,111), (16,954, 4,969, 13,431), and (13,724, 4,637, 32,400), respectively. Scale bar: 500 nm.

In conclusion, our volumetric observations support a simple interpretation: mammalian ERES can consist of vesicle clusters, large vesicles with buds, or extended tubular structures.

## DISCUSSION

We present a unified description of the three-dimensional architecture of ER exit sites (ERES) in mammalian cells, clarifying recent disagreements about the mechanisms underlying the earliest step of the secretory pathway. Specifically, we address the controversy regarding whether tubular or vesicular carriers mediate trafficking from the ER. We obtained high-resolution, isotropic, volume FIB-SEM images from various mammalian cell types preserved by HPFS. These datasets, independently acquired by our laboratory and the COSEM team at Janelia Research Campus, enabled comprehensive three-dimensional reconstructions of this complex subcellular organization. Using automated segmentation powered by our deep-learning-based ASEM pipeline, trained to distinguish COPI vesicles from ER structures, we identified numerous small clusters of 50– 70 nm, COPI-like vesicles closely apposed to flattened ER cisternae. We have termed these clusters, presumably of COPII-coated vesicles, vesicular ERES. Our findings demonstrate that vesicular ERES coexist with tubular ERES -- the latter comprising the tubular extensions previously proposed as exclusive transport carriers originating from the ER.

Our identification of vesicular ERES is consistent with earlier tomographic reconstructions from 250-300 nm thick sections imaged by transmission electron microscopy of cryo-protected samples of yeast (Mogelsvang et al., 2003; Levi et al., 2010) and plants (Donohoe et al., 2013) and of HpG2 cells immuno-gold labeled with an antibody specific for Sec23 and Sec 31 from COPII (Zeuschner et al., 2006). These observations also agree with the 3D architecture of vesicular ERES previously described as clusters of small vesicles adjacent to enlarged ER cisternae in fat cells and imaginal discs from flies imaged using volume FIB-SEM (Yang et al., 2021). In that study, COPII-containing ER buds were shown adjacent to vesicles similarly decorated with COPII. Further support comes from cryogenic high-resolution electron tomography of INS-1 cells, imaged without fixation or staining, which revealed clear examples of coated ER buds adjacent to vesicular clusters containing both coated and uncoated vesicles (Carter et al., 2020). Although the structural resolution in these reconstructions was insufficient to definitively assign the coat composition, the arrangement is consistent with vesicular ERES.

Although broadly accepted, this model has been challenged by the alternative view that COPII facilitates cargo sorting at ERES but does not directly form transport vesicles (Mironov et al., 2003). Rather, COPII may remain at the ER membrane, not associating with cargo carriers moving toward the Golgi apparatus. Support for this hypothesis comes from live-cell fluorescence microscopy experiments showing fluorescently tagged COPII markers remaining stationary at ERES, while cargo-containing structures lacking COPII fluorescence move (Westrate et al., 2020; Hammond and Glick, 2000; Shomron et al., 2021). Furthermore, fluorescence recovery after photobleaching (FRAP) experiments indicated rapid exchange of COPII subunits between ERES- associated spots and the cytosol (Forster et al., 2006). It has also been suggested that stable COPII assemblies at ERES form tubular necks at the base of ER protrusions, which after membrane fission yield COPII-free tubular carriers traveling to the Golgi (Shomron et al., 2021). However, this interpretation faces limitations inherent to the resolution of fluorescence microscopy. The observed fluorescent spots were diffraction-limited (∼300 nm in the x/y plane, ∼600 nm along the z-axis), dimensions consistent with the optical point spread function. Consequently, these images could not reliably discriminate between COPII-coated buds or tubular necks and individual vesicles or vesicle clusters separated by distances of 300–400 nm, as demonstrated by higher-resolution electron microscopy from our studies and those of others (Carter et al., 2020; Zeuschner et al., 2006; Yang et al., 2021). Thus, these experiments do not unequivocally distinguish whether the observed COPII dynamics reflect exchange between membrane-bound and cytosolic pools or represent rapid, asynchronous cycles of vesicle coating and uncoating within unresolved clusters.

To address these discrepancies, (Weigel et al., 2021) combined cryo-structured illumination microscopy (cryo- SIM) with high-resolution FIB-SEM imaging to morphologically characterize ERES, identified by fluorescently tagged COPII subunits and secretory cargo. Their FIB-SEM analysis, primarily conducted at 8 × 8 × 8 nm resolution on HeLa cells overexpressing the fluorescent markers, concluded that tubular extensions alone defined ERES, excluding vesicular structures. However, reliance on colocalization with fluorescent COPII puncta could have inadvertently overlooked detection of vesicular ERES.

Our analysis using higher-resolution FIB-SEM images (5 × 5 × 5 nm and denoised 2 x 2 x 1 nm) clearly revealed vesicular ERES in diverse mammalian cell types. Positional correlation analyses between tubular ERES identified with the ERES model by the COSEM team and vesicular ERES we detected in non-expressing HeLa cells confirm the coexistence of these distinct morphological entities. In alignment with earlier proposals, we propose that vesicular ERES primarily mediate the transport of relatively small cargo, whereas tubular ERES facilitate trafficking of larger cargo. Moreover, the morphological similarities between tubular ERES and vesicular-tubular clusters (VTCs), which correspond to the ER-Golgi intermediate compartment (ERGIC) (Nakano, 2022; Martínez-Menárguez et al., 1999), suggest that some tubular ERES might serve as sorting stations involved in both retrograde and anterograde traffic. Whether the different forms of ERES reflect distinct temporal stages in their biogenesis, or whether the ratio of tubular to vesicular ERES adjusts dynamically in response to increased cargo flux (Forster et al., 2006; Farhan et al., 2008) or to changes in cellular physiology, remains to be determined.

In conclusion, the most parsimonious model integrating the morphological and cell biochemical available data suggests that COPII assembles at specialized ER sites distributed throughout the cell, coordinating cargo capture and sorting with membrane deformation. This assembly process can result in either fully enclosed COPII-coated vesicles or partially assembled coats functioning as scaffolds for extended membrane carriers, such as the tubular structures observed in mammalian cells. Following vesicle formation, COPII rapidly disassembles, often before the vesicle has moved far from its ER membrane of origin, like the uncoating dynamics of endocytic clathrin-coated vesicles (CCVs) and of CCVs originating from the trans-Golgi network. The uncoated COPII vesicles and tubular carriers then traffic toward the Golgi apparatus along microtubules, guided by motor proteins.

Finally, our study highlights how high-resolution electron microscopy, applied at the cellular-volume scale, enables detailed structural insights from 3D imaging data. Our ASEM deep learning pipeline enabled accurate and efficient segmentation of vesicles and ER structures, used here as a tool to guide and facilitate visualization of the inherently complex subcellular architecture in its native spatial context. Importantly, our goal was not exhaustive identification of every vesicle, but rather to provide a reliable visual framework to know “*where to look”*. We provide unrestricted access to our datasets and user-friendly tools developed for the ASEM pipeline. Our publicly available repository, https://open.quiltdata.com/b/asem-project, includes FIB-SEM volumes, training datasets, segmentation models and reconstructions, and our GitHub repository at https://github.com/kirchhausenlab/incasem easily accessible open-source code. By making these resources freely available, we aim to enable the broader scientific community to explore, validate, and extend our findings, fostering further discoveries at the intersection of computational and cell biology.

## Material and Methods

### Cells

Growth conditions for mycoplasma-free human fetal immortalized astrocyte SVG-A cells and iPSC-derived neurons (iNs) were as described previously (Gallusser et al., 2022) (Sanyal et al., 2024), (Gallusser et (Table I). Inmortalized MuTuDC dendritic cells (Pigni et al., 2018) were a gift from Hidde Ploegh laboratory; they were cultured in RPMI media (Corning 10-040-CM) supplemented with 8% FBS, (Atlanta Biologicals S11150H), 50uM of B-mercaptoethanol and incubated with 1μg/ml of lipopolysaccharide(LPS) prior to FIB-SEM imaging. Conditions for HeLa cells were described in (Weigel et al., 2021).

**TABLE I.**
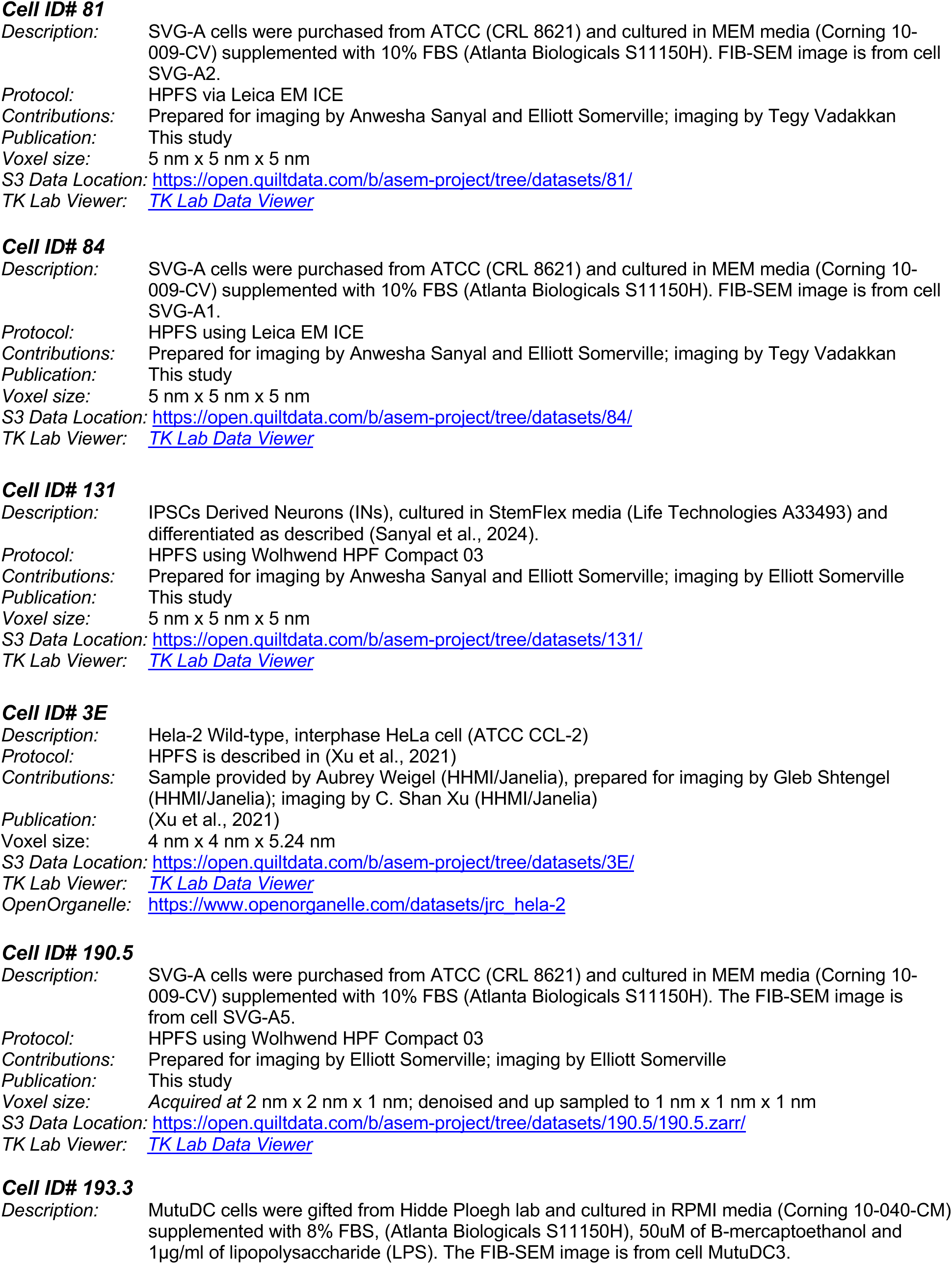
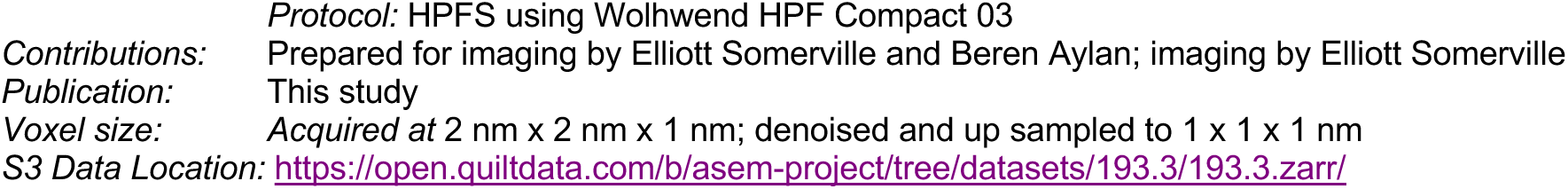

### High-Pressure Freezing, Freeze-Substitution and Embedding

Procedures for high-pressure freezing, freeze-substitution (HP-FS), and embedding of SVG-A cells and iNs were as described previously (Gallusser et al., 2022; Sanyal et al., 2024). Briefly, SVG-A cells were plated on 6 × 0.1 mm sapphire disks (616-100; Technotrade International; six disks per dish) and grown in MEM medium (Corning 10009CV) supplemented with 10% fetal bovine serum (Atlanta Biologicals S11150) (Sanyal et al., 2024) . iNs were plated on 6 x 0.1 mm sapphire disks pre-coated with Matrigel as detailed in (Sanyal et al., 2024). For freezing, two sapphire disks—one with cells—were separated by a 100 µm stainless steel spacer (Technotrade 1257-100) and processed using either a Leica EM ICE (Leica Microsystems; SVG-A cells) or Wohlwend HPF Compact 03 (Technotrade; iNs) high-pressure freezer. Frozen samples were transferred under liquid nitrogen to cryotubes containing frozen substitution medium (2% OsO₄, 0.1% uranyl acetate, and 3% water in acetone).

Freeze-substitution was performed in an EM AFS2 apparatus (Leica Microsystems) programmed for sequential warming: −140°C to −90°C (2 h), −90°C for 24 h, −90°C to 0°C (12 h), and 0°C to 22°C (1 h). Samples on sapphire disks were sequentially rinsed three times with anhydrous acetone, propylene oxide (Electron Microscopy Sciences), and 50% resin (Embed 812 mix; EMS 14121) in propylene oxide. Disks were then transferred into embedding molds (EMS 70900) containing 100% resin and polymerized at 65°C for 48 h. After polymerization, sapphire disks were removed from the resin blocks by sequential immersion in liquid nitrogen and boiling water.

### Crossbeam FIB-SEM Isotropic Imaging

As previously described (Gallusser et al., 2022), polymerized resin blocks were removed from molds and mounted onto aluminum pin stubs (Ted Pella) with conductive silver epoxy (EPO-TEK H20S, EMS), exposing the free face. The exposed face was carbon-coated (20 nm) using a Quorum Q150R ES sputter coater (Quorum Technologies) and loaded into a Zeiss Crossbeam 540 microscope for FIB-SEM imaging. After eucentric correction, the stage was tilted to 54° with a working distance of 5 mm for beam alignment. Following SEM localization of the cell of interest, a platinum protective layer was deposited by gas injection. A coarse trench adjacent to the cell was milled with a 30 kV/30 nA gallium ion beam, and the exposed block face was polished at 30 kV/7 nA.

Sequential imaging employed interlaced milling (30 kV/3 nA gallium beam) and SEM imaging (1.5 kV/400 pA) at 5 nm intervals, yielding isotropic voxels (5 nm in X/Y/Z) or SEM imaging (1.5 kV/800 pA) at 1 nm intervals for anisotropic voxels (2x2x1 nm in X/Y/Z). Registration marks were created atop the platinum layer with a 1.5 kV/50 pA gallium beam, enhanced by 1.5 kV/5 nA electron irradiation, followed by a secondary platinum coating. FIB-SEM images captured at 5 x 5 x 5 or 2 x 2 x 1 nm represented averaged signals from the Inlens secondary electron and backscattered electron ESB detectors, captured with a pixel dwell time of 10–15 μs or 1-2 μs, respectively. Sequential z-plane images were aligned using the Fiji plugin Register Virtual Stack Slices (https://imagej.net/plugins/register-virtual-stack-slices) with translation and shrinkage constraints (Schroeder et al., 2021).

### Denoising of FIB-SEM images

We implemented a denoising strategy we named FastFIB, a self-supervised denoising approach based on the Noise2Noise paradigm (Lehtinen et al., 2018), which trains a neural network to predict one noisy measurement from another, assuming both share an underlying signal. Because the network cannot reconstruct the uncorrelated noise, it learns to suppress it, yielding a cleaner output. In our case, the training pairs consisted of corresponding 3D sub volumes from FIB-SEM datasets acquired with inlens and ESB detectors.

FastFIB uses a 3D U-Net architecture with four levels of depth and approximately 10 million trainable parameters. The loss function was the Charbonnier loss, defined with α = 0.5 and ε = 1 × 10⁻³. Prior to computing the loss, we applied a mild 3D Gaussian blur to the U-Net output. This procedure encouraged the network to produce slightly deblurred outputs, as the blurred version had to match the target. To further regularize the output, we penalized the spatial gradients of the difference between the raw output and its blurred counterpart, promoting smoothness and suppressing high-frequency noise.

We trained the model on two datasets: one we acquired at isotropic 5 × 5 × 5 nm resolution, and the other at anisotropic 2 × 2 × 1 nm resolution. Before training, we upsampled the latter to 1 × 1 × 1 nm using trilinear interpolation to enforce isotropy.

After training, we performed brief finetuning on our FIB-SEM data acquired 5 × 5 × 5 nm or 2 × 2 × 1 nm data that had not been used for training, which modestly improved the network’s performance on these inputs. We also applied the trained 5 × 5 × 5 nm FastFIB model directly to the 4 × 4 × 5.24 nm FIB-SEM dataset of a HeLa cell from the COSEM project. Because this dataset included only inlens detector data, no retraining was performed.

### Annotations

#### COPI Vesicles

Ground truth manual annotations for selected COPI vesicles (50–70 nm diameter) adjacent to the Golgi apparatus in SVG-A cells were performed approximately every third plane, followed by automatic volumetric completion using WebKnossos (Boergens et al., 2017). Binary masks were created at least 47 voxels from the edges of the 3D FIB-SEM images from three regions adjacent to the trans-Golgi network in SVG-A1 cells, containing 42, 47, and 51 vesicles, respectively, and one region in SVG-A2 cells with 32 vesicles. Dimensions of the training regions were 204 x 204 x 204, 110 x 250 x 250 and 225 x 225 x 225 voxels in SVG-A1 and 225 x 225 x 225 voxels in SVG-A2; dimensions of the validation region in SVG-A1 were 180 x 180 x 210 voxels.

#### Dispersed COPI–like Vesicles and Vesicular ERES

The COPI ASEM model trained with the selected COPI vesicles identified numerous vesicles in naïve regions of the SVG-A cells not used for training. To catalog these, voxels labeled as positive within 60 nm were grouped into single objects (e.g., 331 vesicles). Visual inspection revealed many of these as bona fide COPI vesicles clustered near the Golgi apparatus (Fig. 2). Vesicles in regions larger than 400 nm or within 320 nm in any dimension were computationally excluded, leaving 273 dispersed vesicles throughout the cell. Of these, visual inspection of the 247 identified objects were individually or occasionally paired vesicles clustered with similar vesicles undetected by the COPI ASEM model.

We calculated the shortest geometric distance between these vesicles and ER structures segmented using our ER ASEM model, identifying 247 vesicles within 120 nm of the ER, designated vesicular endoplasmic reticulum exit sites (vesicular ERES). We calculated the distance between the vesicle membrane and the ER by applying the Pythagorean theorem in three dimensions to coordinates obtained by visual inspection. Neuroglancer-based visual inspection confirmed that 179 vesicles belonged to clusters of 5–55 vesicles in regions ≤400 nm, while the remaining 68 appeared as isolated vesicles near tubular ERES identified by the COSEM project (Fig. 6).

#### Relationship Between Vesicular and Tubular ERES in the Hela-2 cell

To compare vesicular ERES identified by the COPI ASEM model with tubular ERES defined by the COSEM project in the HeLa-2 cell, FIB-SEM data were analyzed as follows: (1) Voxels labeled as positive by the COSEM ERES model were grouped into single objects if they were within 140 nm of each other (n = 779), a threshold selected based on visual inspection due to larger predicted tubular ERES sizes. (2) X/Y/Z coordinates of 247 vesicular ERES predicted by the COPI ASEM model were compared with coordinates of tubular ERES from the COSEM ERES model to identify vesicles located within 50 nm. (3) Predictions were classified into isolated vesicular ERES (n = 179), vesicular ERES associated with one (n = 68) or two (n = 6) tubular ERES, and isolated tubular ERES (n = 705). Classification accuracy was confirmed by visual inspection using Neuroglancer.

### Computational Requirements for training and predictions

Data fetching and augmentation were executed in parallel using eight CPU cores, and training was performed on a single Nvidia DGX-A100 GPU. Each iteration of the ASEM neural network training required approximately 1 second, with total training durations typically ranging from 100,000 to 170,000 iterations (∼20–30 hours,

including validation). Predictions performed using 100 workers, processing input and output volumes of 364 × 364 × 364 voxels and 270 × 270 × 270 voxels respectively, required approximately 8 minutes per single-cell image stack.

### Model Training and Predictions

#### COPI Vesicles

The ASEM COPI model training using FIB-SEM data from SVG-A1 cells converged after 167,000 iterations, achieving a Dice coefficient of 0.392 and loss function (binary cross entropy) value of 0.008. Visual inspection using Neuroglancer of predicted segmentations (∼4.08 billion cubic voxels), which were not part of the training set, confirmed accurate identification of bona fide COPI vesicles adjacent to the Golgi and COPI-like vesicles distant from the Golgi. These segmentations represented vesicles with diameters of 50–80 nm, alone or as clusters of 2–7. Similar accuracy was observed upon applying the COPI ASEM model to iN and HeLa cell datasets not used during training.

#### Clathrin-Coated Vesicles

Previously, we demonstrated the efficacy of the clathrin-coated pit ASEM model trained on endocytic clathrin-coated pits for identifying clathrin-coated vesicles near the cell surface and distal side of the trans-Golgi network (TGN) (Table I, cells 12 and 13 (Gallusser et al., 2022)). Using this model, we predicted clathrin-coated vesicles in the SVG-A1 cell. Neuroglancer-based visual inspection verified these predictions, identifying vesicles near the TGN ranging in diameter from 110 to 130 nm and clearly distinct from COPI vesicles detected by the COPI ASEM model.

### Statistical Analysis

The FIB-SEM samples from different cell types are biological replicates, e.g. fully processed independently of each other. Because the study focuses on description of non-quantifiable morphological features, statistical testing is not applicable and was not applied.

## Data availability

The datasets of raw FIB-SEM cell images, ground truth annotations, probability maps predicted by the models, and corresponding segmentation masks are publicly available at the AWS ASEM bucket: https://open.quiltdata.com/b/asem-project.

An online tool has been created to view 3D volumes relevant to this study and others. The tool is available at http://asem-viewer-env.eba-rrnvmfwa.us-east-1.elasticbeanstalk.com/ and allows users to quickly select and visualize cells and volumes (raw data, prediction results, and ground truth labels) in the browser using Neuroglancer.

## Code Availability & Usage

The ASEM pipeline software and step-by-step usage instructions are publicly available at https://github.com/kirchhausenlab/incasem (ASEM deep learning pipeline). Users comfortable with Linux and who have access to an NVIDIA GPU should follow the standard installation and usage instructions from GitHub.

For users with limited experience in these areas or without a GPU, we provide two alternatives to facilitate usage. <u>Option 1:</u> A Google Colab notebook, available as ASEM_Notebook.ipynb, provides a cloud-based alternative that eliminates the need for a local installation. The notebook can be access from the *Interactive Demo* section of the README.md in the incasem GitHub. This option is ideal for users who wish to explore ASEM as a demo and access cloud-hosted GPUs. Note that a Colab membership may be required for extended use.

<u>Option 2:</u> Through the GitHub installation, users can run incasem in command-line mode or via a GUI that assists with configuring and launching training, fine-tuning, and predictions. For access details, see *the UI Installation*section of README.md in the incasem GitHub.

Trained neural network models are available at https://open.quiltdata.com/b/asem-project, with usage instructions provided at https://github.com/kirchhausenlab/incasem. The models used to detect COPI vesicles for the primary analysis is 2631.

**Video 1.** Manual annotations of COPI vesicles (blue) adjacent to the Golgi apparatus, rendered through the passing FIB-SEM volume of the same SVG-A cell shown in Fig. 2A and B, using WebKnossos.

**Video 2.** Manual annotations of COPI (blue) and clathrin-coated vesicles (cyan) adjacent to the Golgi apparatus, rendered through the FIB-SEM volume of the SVG-A cell shown in Fig. 2C, using WebKnossos.

**Video 3.** Manual annotations of COPI vesicles (blue) adjacent to the Golgi apparatus, rendered through the passing FIB-SEM volume of the iN cell shown in Fig. 2D, using WebKnossos.

**Video 4.** Manual annotations of COPI vesicles (blue) adjacent to the Golgi apparatus, rendered through the passing FIB-SEM volume of the HeLa cell shown in Fig. 2E, using WebKnossos.

**Video 5.** Manual annotations of COPII vesicles (blue) adjacent to the annotated ER (yellow), identified as vesicular ERES rendered through the passing FIB-SEM volume of the SVG-A cell shown in Fig. 3A, using WebKnossos.

**Video 6.** Manual annotations of COPII vesicles (blue) adjacent to the annotated ER (yellow), identified as vesicular ERES rendered through the passing FIB-SEM volume of the iN cell shown in Fig. 3B, using WebKnossos.

**Video 7.** Manual annotations of COPII vesicles (blue) adjacent to the annotated ER (yellow), identified as vesicular ERES rendered through the passing FIB-SEM volume of the HeLa cell shown in Fig. 3C, using WebKnossos.

**Video 8**. Manual annotations of COPI vesicles (blue) adjacent to the Golgi apparatus (green) rendered through the passing FIB-SEM volume of the SVG-A cell shown in Fig. 4. Annotations were performed using the Labkit plugin in FIJI (Arzt et al., 2022), and masks were exported to Imaris (Bitplane, Zurich, Switzerland) for 3D rendering using the “Surfaces” feature. The movie was generated with the “Animation” tool in Imaris.

**Video 9.** Manual annotations of COPII vesicles (blue) adjacent to the annotated ER (yellow), identified as vesicular ERES rendered through the passing FIB-SEM volume of the MuTuDC cell shown in Fig. 5.

Annotations were performed using the Labkit plugin in FIJI (Arzt et al., 2022), and masks were exported to Imaris (Bitplane, Zurich, Switzerland) for 3D rendering using the “Surfaces” feature. The movie was generated with the “Animation” tool in Imaris.

**Video 10.** Manual annotation of COPII vesicles (blue) adjacent to the annotated ER (yellow), identified as vesicular ERES rendered through the passing FIB-SEM volume of a region of the HeLa cell shown in Fig. 6A, using WebKnossos.

**Video 11.** Manual annotation of a tubular ERES (dark orange) adjacent to the annotated ER (yellow), rendered through the passing FIB-SEM volume of a different region of the HeLa cell shown in Fig. 6A, using WebKnossos.

**Video 12.** Example of a mixed ERES, illustrated by manual annotations in WebKnossos. A single COPII-coated vesicle (blue), adjacent to the annotated ER (yellow), defines a vesicular ERES in close spatial association with a tubular ERES (dark orange). The annotations are rendered through the FIB-SEM volume of a distinct region of the HeLa cell shown in Fig. 6A.

## ACKNOWLEDGMENTS

We thank J. O’Connor for his help setting up and maintaining the computing infrastructure and to members of the Kirchhausen laboratory for help and encouragement. The research was supported by a National Institute of General Medical Sciences Maximizing Investigators’ Research Award GM130386 and by the NNF Center of Optimized Oligo Escape and Control of Disease awarded to T. Kirchhausen and N. Hatzakis. Athul Nair and Akhil Nair were supported in part by discretionary funds available to T. Kirchhausen. Acquisition of the FIB-SEM microscope was supported by a generous grant from Biogen to T. Kirchhausen, and the high-pressure freeze substitution device was made available by S.C. Harrison. Acquisition of the computing hardware including the DGX’s GPU-based computers, CPU clusters, fast access memory, archival servers, and workstations that made possible this study were supported by generous grants from the Massachusetts Life Sciences Center to T. Kirchhausen and by an equipment supplement to the National Institute of General Medical Sciences Maximizing Investigators’ Research Award GM130386. Construction of the server room housing the computing hardware was made possible with generous support from the PCMM Program at Boston Children’s Hospital.

The authors declare no competing financial interests.

## Author contributions

T. Kirchhausen conceptualized and designed the experiments, drafted the manuscript, and finalized it in close consultation with all co-authors. A. Nair and A. Nair, as undergraduate students, generated and curated the ground truth annotations used for training, implemented the computational pipeline to train the ASEM neural network, obtained models, made predictions, and analyzed the volumetric FIB-SEM data. E. Somerville and A. Sanyal prepared the samples for FIB-SEM. E. Somerville also maintained the FIB-SEM instrument and performed data collection and pre-processing. J.I. Costa-Filho developed FastFIB, the neural network–based tool for denoising volumetric FIB-SEM data, which enabled acquisition of high-resolution datasets. P. Stock developed the online tool for 3D data visualization, created the publicly accessible AWS data repository and supported deep learning and data analysis efforts. A. Jain implemented our publicly available ASEM user interface software and usage instructions.

